# Surveying Tropical Faunal Diversity via Airborne DNA Analyses

**DOI:** 10.1101/2025.07.25.666788

**Authors:** Joseph M. Craine, Nicholas Schulte, Jessica Devitt, Devin Leopold, Kristin Saltonstall, Noah Fierer

## Abstract

Here, we assess the potential of airborne eDNA (airDNA) metabarcoding to characterize arthropod and vertebrate assemblages in a tropical forest. We deployed 28 high-flow air samplers across a 1.5 km^2^ area on Barro Colorado Island, Panama, over two consecutive 24- hour periods and sequenced captured DNA using general eukaryotic and vertebrate-specific metabarcoding primers. AirDNA sampling detected 1293 arthropod operational taxonomic units (OTUs) alongside 157 vertebrate OTUs representing birds, mammals, reptiles, amphibians, and fish. Arthropod richness was comparable to that quantified locally with light and Malaise trap surveys, while vertebrate richness exceeded that typically observed with conventional techniques. The level of field and lab effort employed in this study captured approximately 84% of the asymptotic richness for arthropods and 76% for vertebrates, implying that relatively little additional richness would have been quantified with additional effort within the same domain of sampling. Observed and asymptotic richness increased with the number of sites, the number of days a site was sampled, and the number of PCR replicates run per sample, highlighting the need to standardize effort among future applications of airDNA to compare richness across systems. These results demonstrate that airDNA metabarcoding can efficiently and non-invasively profile tropical terrestrial faunal biodiversity.

## Introduction

As biodiversity continues to decline globally, there is an urgent need for effective monitoring tools to inform conservation efforts and sustainable land management^1,2^. Biodiversity monitoring aims to assess the presence, abundance, and distribution of species across time and space^3^. However, effectively quantifying the diversity of terrestrial animal assemblages across diverse habitats remains a major challenge^4^.

Sampling environmental DNA (eDNA) has the potential to fundamentally alter terrestrial biodiversity monitoring by providing a broad zone of non-invasive monitoring with species-specific identification. The term eDNA refers to genetic material shed by organisms into their surrounding environment, but also includes the DNA in small-bodied organisms and any prey that has been consumed by organisms that are collected simultaneously^5^. This DNA can be collected from the environment and analyzed using metabarcoding, a technique that amplifies and sequences specific genetic markers, enabling the identification of multiple species from a single DNA sample^6^. Metabarcoding can be used to profile a wide range of taxonomic groups, including vertebrates, insects, plants, fungi, and bacteria from a variety of sample types^7,8^.

Aquatic eDNA metabarcoding is now routinely used for aquatic biodiversity surveys^9,10^, but its utility for capturing terrestrial signals is restricted to niche cases, since terrestrially derived eDNA in adjacent aquatic ecosystems is typically found in concentrations that are too low to reliably index terrestrial biodiversity^11^. Soil holds eDNA from aboveground organisms but faces challenges of DNA degradation and patchy distribution, which complicates reliable inference about terrestrial biodiversity at relevant spatial scales^12^. Other techniques, such as collecting eDNA from surfaces like leaves or from the digestive system of consumers such as leeches or carrion flies can be useful for particular taxa but are likely limited in their ability to capture a broad range of terrestrial animal diversity^13–16^.

Metabarcoding of airborne eDNA, here referred to as airDNA^17^, has the potential to advance terrestrial biodiversity monitoring as much as aquatic eDNA metabarcoding has revolutionized aquatic biodiversity monitoring^18^. Metabarcoding of airDNA can capture DNA from a broad range of aboveground organisms, including terrestrial vertebrates, insects, and plants. This is because many organisms shed DNA into the air, which can then be collected by drawing air through a filter, followed by extraction and metabarcoding of the captured DNA to identify the species present in a given area^19–21^. Recent studies have detected mammalian and avian DNA from air samples collected in both zoo enclosures and nearby natural settings^17,20^. Vertebrate richness from airDNA samples is typically modest, but not markedly lower than conventional techniques. For example, Lynggaard et al.^22^ detected 64 bird, mammal, fish, and amphibian taxa with 144 12-h airDNA samples over three days at 24 forest sites in Denmark – about 25% of known vertebrate diversity at those sites. Insect richness estimates from air samples are typically in the dozens of species per deployment^21^, which is comparable to insect richness recovered by bioacoustics in tropical forests^23^ but much lower than those from conventional malaise or light traps, which can collect hundreds of species in a typical deployment^24,25^.

Given that the concentrations of airborne DNA from terrestrial organisms are extremely low^26^, one bottleneck for routine airDNA metabarcoding is the requirement that a sufficient volume of air is sampled. For instance, cyclonic air samplers typically process less than 4 m³ h^-1^ ^21^, standard Hirst-type volumetric spore traps draw air at ∼0.6 m^3^ h⁻¹ ^27^, while fans used for airDNA capture draw 0.3 – 3.5 m^3^/h through filters^20,28^. Another potential bottleneck might be how much template is processed in PCR reactions per sample. A typical sample might generate 100 µL of DNA extract of which only 1 - 3 µL of template are used in PCR reactions to amplify target DNA. Typically, just one or a few PCR replicates are processed for an eDNA sample, but species richness accumulates with additional numbers of PCR replicates per sample^29^. The marginal benefits of additional PCR replicates for airDNA-based surveys of terrestrial animals are unknown, leaving open questions about how best to maximize diversity capture from airDNA samples.

Despite the promise of airDNA as a survey tool, studies have yet to show that airDNA can index biodiversity at similar levels to conventional assessments in an economical, representative, and informative manner. Here, we test whether airDNA metabarcoding can reconstruct the arthropod and vertebrate assemblages of a well-studied tropical forest by setting up 28 high-flow fans distributed across an area of approximately 1.5 km^2^ and sampling airDNA over two consecutive 24-hour periods. DNA was extracted from filters and amplified with both a general eukaryotic primer set that would amplify the DNA of vertebrates and arthropods as well as a vertebrate-specific primer set. Twelve PCR replicates were generated and indexed independently for each primer set for each sample. We then used these data to quantify rates of species accumulation as a function of laboratory and field effort to begin to understand the utility of monitoring terrestrial animal biodiversity via airDNA analyses and how to do so most efficiently.

## Results

### Arthropod diversity assessed via airDNA analyses of COI

Sequencing the 56 samples with 12 independent PCR replicates per sample for the COI primers generated 1293 arthropod and 14 vertebrate OTUs at 97% similarity (Table S2, Text S1). For the arthropods, the mean percent similarity of OTUs to the top NCBI or BOLD reference hit was 95.7% (Table S3). 103 OTUs matched at 100% to a sequence in the reference library and 267 at 99% or higher.

Of the 1293 arthropod OTUs, 1052 OTUs (81%) were assigned to Diptera (flies), and 982 (93%) of Diptera OTUs were assigned to Cecidomyiidae (gall midges) (Figure 1a). Therefore, 76% of all arthropod OTUs were assigned to Cecidomyiidae, and 7 of the 10 most abundant OTUs matched to taxa in Cecidomyiidae family, with the most abundant OTU found in 490 of the 672 PCR replicates. Of the 70 non-Cecidomyiidae Diptera OTUs, 47 were assigned to a family, with Psychodidae (20), Phoridae (11), and Chironomidae (10) being the most abundant (Figure 1).

**Figure 1.**
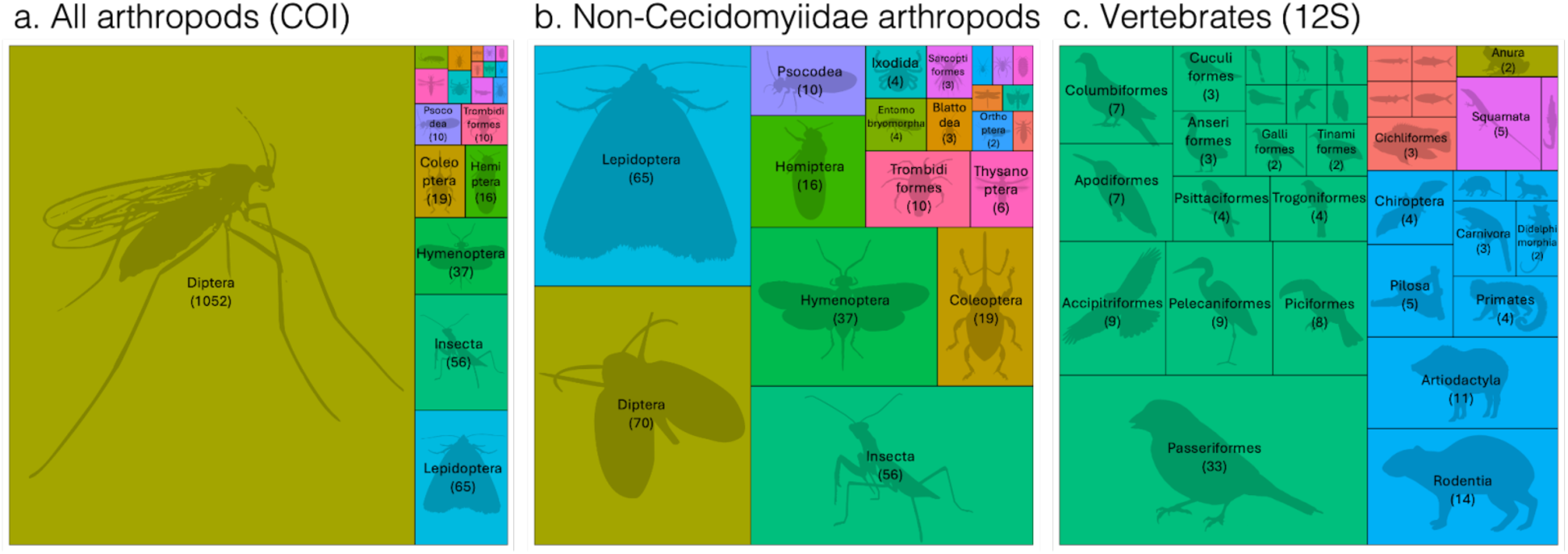
Number of OTUs by order for (a) all COI arthropods, (b) all COI arthropods excluding Cecidomyiidae, and (c) all 12S vertebrates. Unlabeled orders had < 10 OTUs each for (a) and 1 OTU each for (b) and (c).

Other abundant arthropod orders were Lepidoptera (65 OTUs), Hymenoptera (37), Coleoptera (19), Hemiptera (16), Trombidiformes (10), and Psocodea (10) (1b). Of the 65 OTUs assigned to Lepidoptera, 44 were assigned to 15 different families, with Geometridae (12) and Pyralidae (6) having the highest richness. Twenty-one Lepidoptera OTUs were not assigned unambiguously to a single family.

When summed across PCR replicates, arthropod OTU richness per sample varied from 16 to 316 (Table S1). As the number of PCR replicates per sample increased, the number of arthropod OTUs increased in an asymptotic fashion (Figure 2a). With 12 PCR replicates, the mean number of arthropod OTUs per sample was 141. If we doubled the number of PCR reps, we estimate that only 21 more arthropod OTUs would be added per sample on average (Figure 2a).

**Figure 2.**
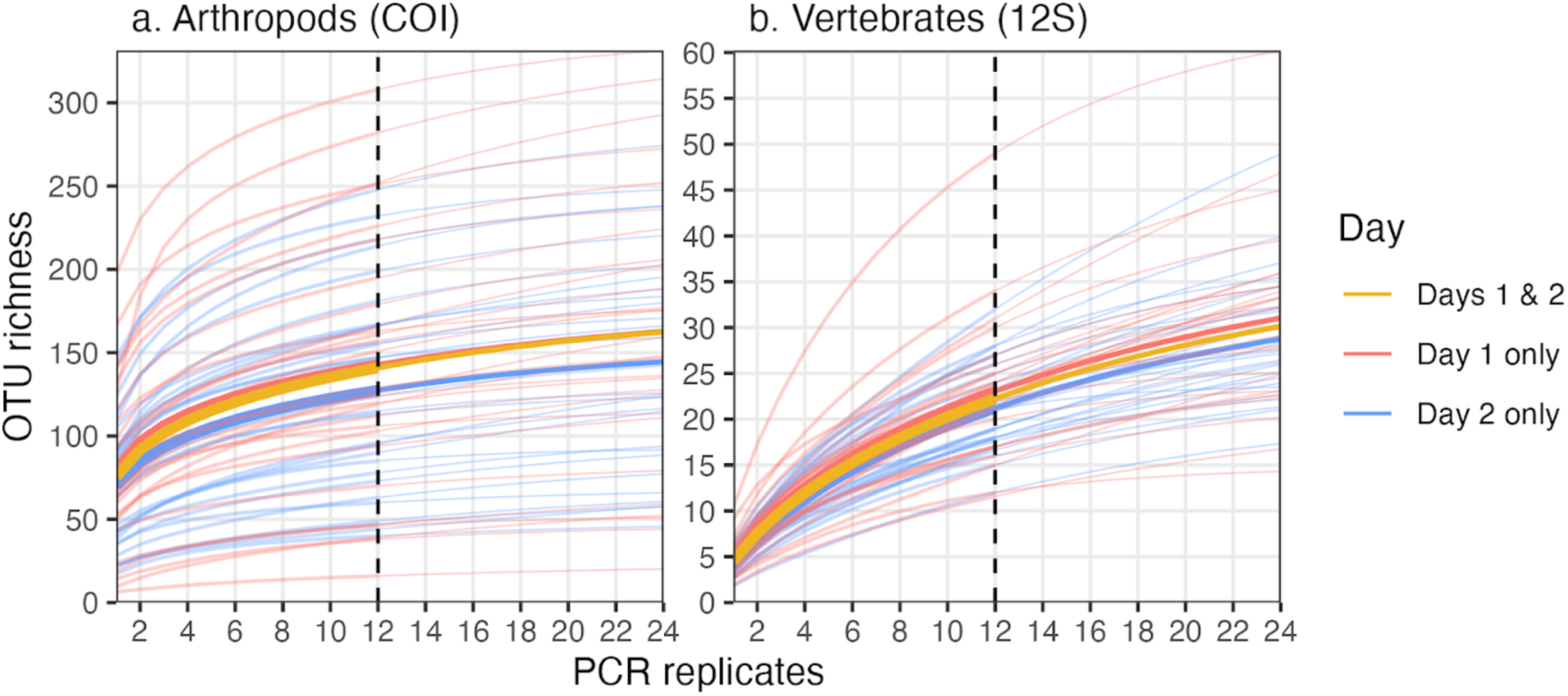
Accumulation of (a) COI arthropod and (b) 12S vertebrate OTUs across PCR replicates for each sample (thin lines) and the mean across samples (bold lines). Averages shown for samples taken on Day 1, Day 2, and the two days combined per site. Vertical dashed lines represent the number of PCR replicates run per sample (n = 12), and thin segments of curves represent extrapolated accumulation to double the number of PCR replicates.

Across the 28 sites, the number of arthropod OTUs increased with additional sites, with 1293 arthropod OTUs from the 28 sites capturing 84% of the asymptotic richness (1549 OTUs) (Figure 3a). On average, a single site generated 295 arthropod OTUs and a second site added an additional 151 arthropod OTUs. If the number of sites were to be doubled from 28 to 56, richness is projected to increase by 15% to 1481 arthropod OTUs (Figure 3a).

**Figure 3.**
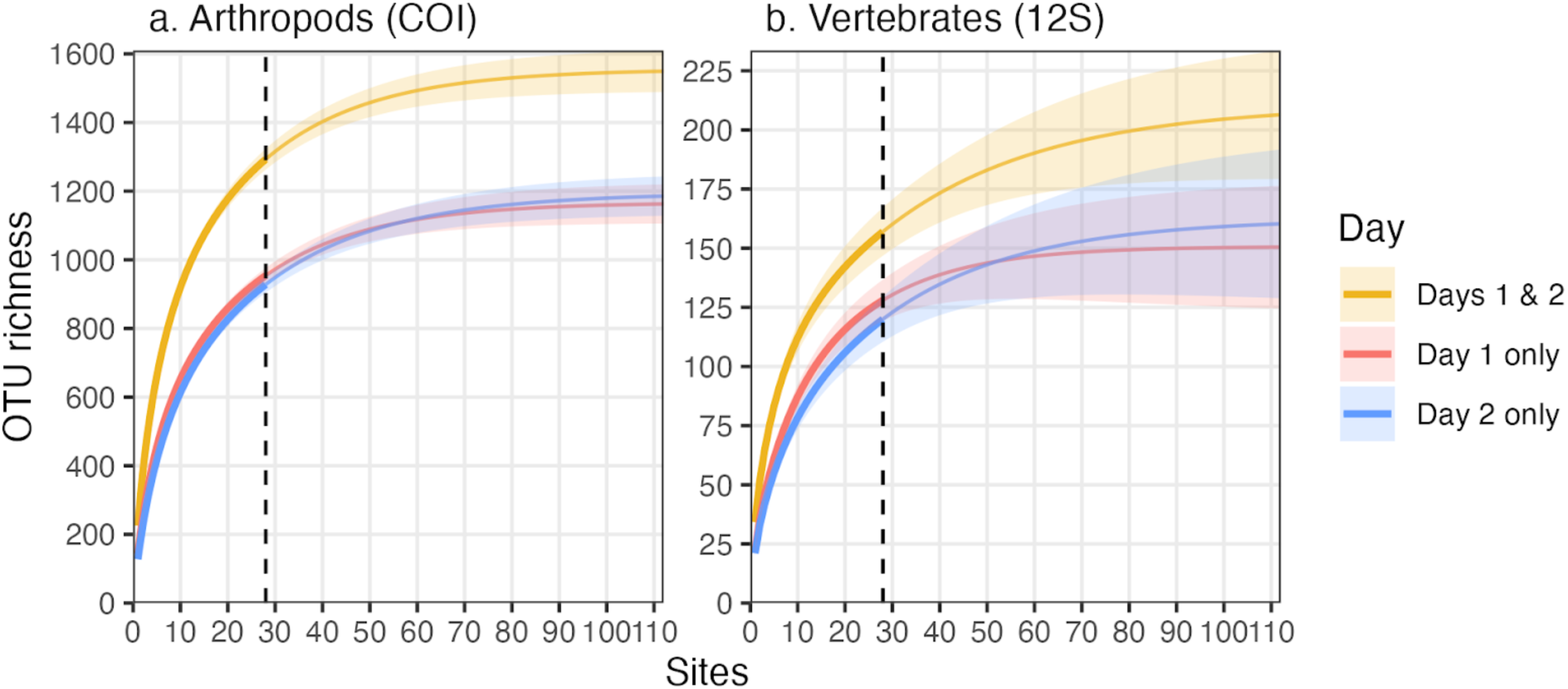
Accumulation of OTUs by adding sampling sites for (a) arthropods and (b) vertebrates across samples after summing PCR replicate reads per sample. For Days 1 & 2, OTU reads were summed by site across days. Vertical dashed lines represent the number of sites (n = 28), and thin segments of curves represent extrapolated accumulation to 4x the number of sites.

Proximity between sites did not correspond to more similar assemblage composition: there was no significant relationship between geographic distance and pairwise similarity in arthropod assemblage composition (Mantel Spearman *p* > 0.47 for both Jaccard and Bray-Curtis distances) (Figure S1a,b).

There was good correspondence among sites in the arthropod OTU richness observed between the two days: sites that had more species on day 1 had more species on day 2 (RMA R^2^ = 0.59, p < 0.001) (Figure S2a). Sites that had higher richness on day 1 were associated with a greater number of unique taxa on day 2. This marginal increase was just 20 unique OTUs on day 2 for every additional 100 OTUs on day 1 (Figure S2b). Repeated sampling of the same site on consecutive days increased richness less than adding an additional site (permutation test *p* < 0.001, Figure 4). For a given site, the second day added 53 OTUs on average (Figure S2b), which was 49% of the marginal richness added if the second sample from a different location was added on the first day (104 OTUs) (Figure 3a).

**Figure 4.**
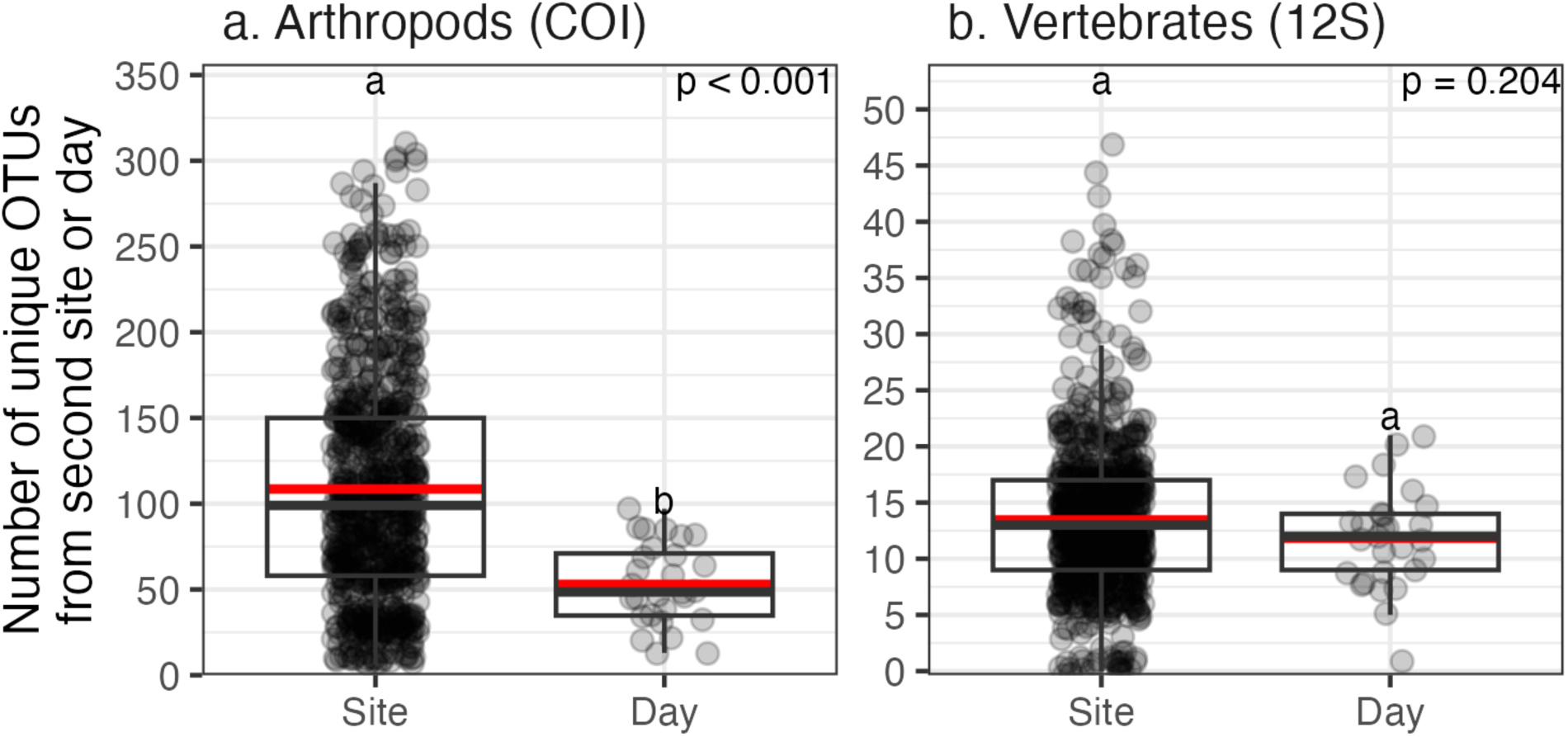
Marginal (a) COI arthropod and (b) 12S vertebrate OTU richness between sites on day 1 and between days for each site. Red lines represent the means, and p-values and letters reflect results from permutation tests.

Exploring the controls on predicted asymptotic richness for arthropods at the site level, decreasing the number of independent PCR replicates summed for a single sample led to lower observed richness and lower predicted asymptotic richness estimates (Figure 5). At 1 PCR replicate per sample, arthropod richness summed across all 56 samples would have averaged 22% less than with all 12 PCR replicates (1000 vs. 1293 OTUs) (Figure 5a). Asymptotic richness with 1 PCR replicate per sample would have been 12% lower than that with 12 PCR replicates per sample, with an estimated asymptotic richness of 1357 vs. 1538 OTUs (Figure 5b). When estimating the effect of doubling sequence depth across all samples, richness was predicted to increase by just 3 OTUs (1296 OTUs vs. 1293 observed). Running the fans for just 1 day instead of adding a second day of samples generated 26% fewer arthropod OTUs (954 for day 1 vs. 1293 for both days) and an asymptotic richness that was 25% lower (1163 for day 1 vs. 1549 for both days) (Figure 3). On average, if we had sampled only half as many site locations (i.e. 14 sites), observed arthropod richness would have been 43% lower with day 1 only and 18% lower across both days than sampling the full 28 sites both days, while asymptotic richness would have been 38% lower with day 1 only (844 vs. 1163) and 33% lower with both days (1164 vs. 1549) (Figure S3).

**Figure 5.**
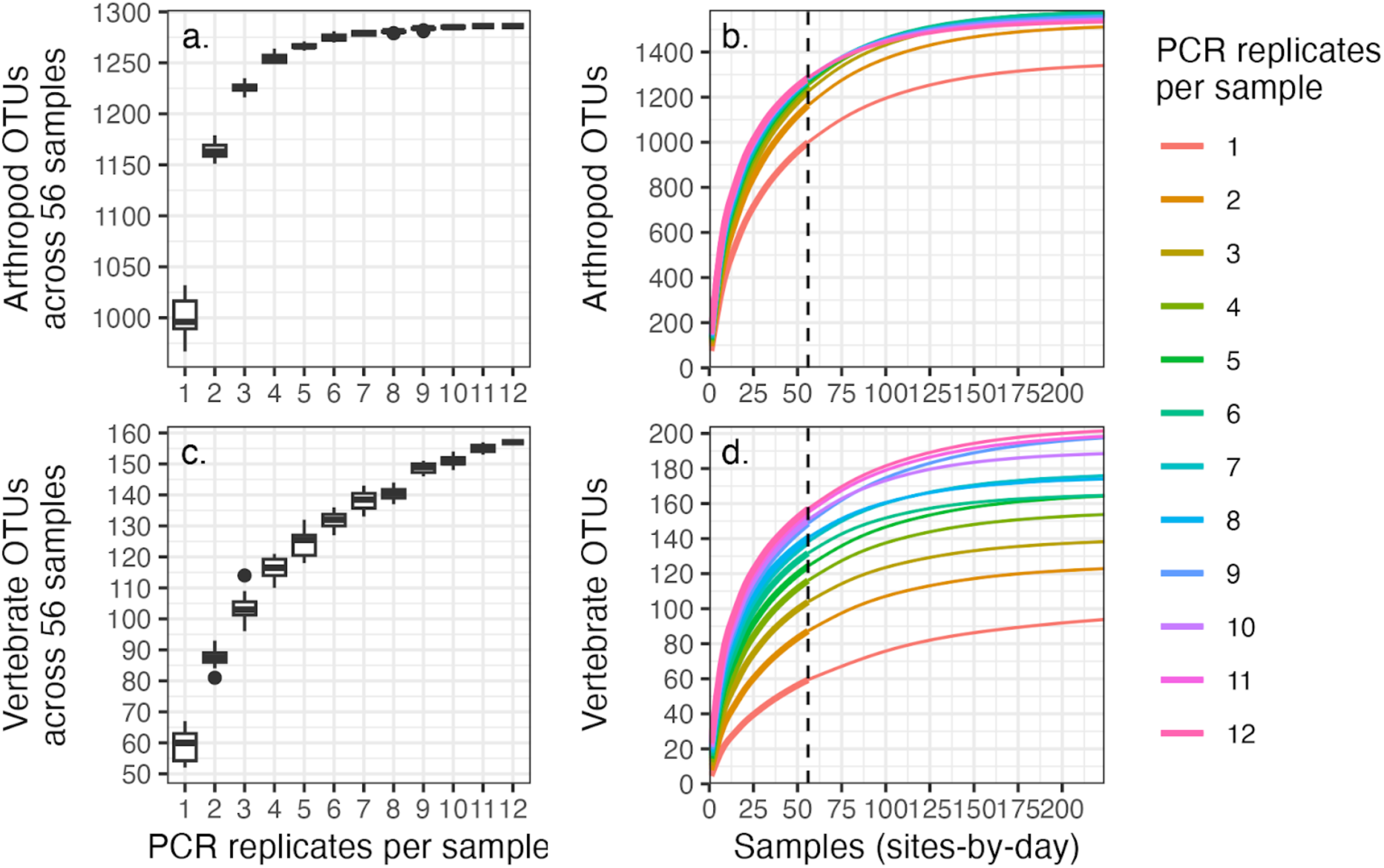
Distribution of (a,b) COI arthropod and (c,d) 12S vertebrate OTU richness across all samples when varying the number of independent PCR replicates per sample. (a,c) Observed richness across 56 samples with up to 99 random draws of the PCR replicates per number of PCR replicates per sample. (b,d) Observed and extrapolated OTU accumulation to asymptotic richness based on up to 99 random draws of PCR replicates per number of PCR replicates per sample (air filter from one site on one day). Vertical dashed lines represent the number of samples (n = 56), and thin segments of curves represent extrapolated accumulation to 4x the number of samples.

### Vertebrate diversity assessed via airDNA analyses with 12S rRNA

Sequencing the 56 samples with 12 independent PCR replicates per sample for the 12S rRNA primers generated 157 OTUs at 97% similarity (Table S4). Of the 157 OTUs, the mean percent similarity of OTUs to the top reference hit was 97.2% (Table S5). 43 of these OTUs matched at 100% to a sequence in the reference library and 74 at 99% or higher. Besides peccary (10 OTUs; only 2 species recognized in Panama), there was no obvious inflation of species counts from the OTUs for any taxonomic group (Table S4).

Of the 157 vertebrate 12S rRNA OTUs, 97 were assigned to Aves, 45 to Mammalia, 7 to Actinopteri, 6 to Reptilia, and 2 to Amphibia (Figure 1c). The 97 bird OTUs were distributed among 18 orders, with Passeriformes the richest (33 OTUs) followed by Accipitriformes (9), Pelecaniformes (9), and Piciformes (8) (Figure 1c). The bird OTUs that occurred most frequently across replicates include *Coragyps atratus* (black vulture, 100% match, 135 occurrences out of 672 PCR replicates), *Amazona farinosa* (southern mealy amazon, 99.5%, 122), *Tinamus major* (great tinamou, 98.2%, 105), and *Cardellina canadensis* (Canada warbler, 98.5%, 96).

Mammal OTUs that occurred most frequently across replicates include *Dicotyles tajacu* (collared peccary, 98.7% match, 273 occurrences), *Nasua narica* (white-nosed coati, 95.3%, 229), *Cuniculus paca* (lowland paca, 99.5%, 191), *Dasyprocta punctata* (Central American agouti, 94.9%, 185) *Bradypus pygmaeus / variegatus* (three-toed or brown sloth, 100%, 138). Of the 19 mammal OTUs with unambiguous species assignments, all have been observed on BCI except *Rattus norvegicus* (brown rat), which is present in Panama. Primates included *Alouatta palliata* (mantled howler monkey), an *Ateles* species (spider monkey), a *Cebus* species (capuchin monkey), and a *Lagothrix* species (which is not found on BCI, had just 92.6% reference match, and 1 occurrence). Bats included *Saccopteryx bilineata* (Greater sac-winged bat), *Uroderma bilobatum* (Tent-making bat), *Vampyrodes caraccioli* (Great stripe-faced bat), and a Vespertilionidae (vesper bat) species. All 5 mammal species detected from COI (Text S1) were detected with 12S rRNA. Actinopteri included 3 cichlids (*Cichla*, *Oreochromis*, *Vieja*) as well as marine species such as *Polydactylus virginicus* (100% match). Reptiles include *Crocodylus acutus* (American crocodile), *Boa constrictor* (common boa), an Anolis lizard, two geckos, and *Iguana iguana* (green iguana). Amphibians were identified as *Rhaebo haematiticus* (leaf litter toad) and *Rhinella marina* (cane toad).

When summed across PCR replicates, vertebrate OTU richness per sample varied from 2 to 49. Examining the patterns of vertebrate OTU accumulation, as the number of PCR replicates per sample increased, the number of OTUs increased in an asymptotic fashion (Figure 2b). With 1 PCR replicate, the average number of vertebrate OTUs was 5. With 12 PCR replicates, the average number of vertebrate OTUs was 22. If we had doubled the number of PCR replicates per sample to 24, 8 more vertebrate OTUs were estimated to be added on average to each sample.

Examining patterns of OTU accumulation across the 28 sites, the number of OTUs increased strongly with additional sites (samples from days 1 and 2 combined), with 28 samples capturing 156 OTUs, which was 76% of the asymptotic richness of 206 OTUs (Figure 3b). On average, one site generated 34 OTUs and the next site an additional 17 OTUs. If the number of sites were to be doubled from 28 to 56, vertebrate richness was estimated to increase by 31% (Figure 3b). Proximity between sites did not correspond to more similar community composition: there was no significant relationship between geographic distance and pairwise similarity in OTU presence/absence or relative abundance (Mantel Spearman *p* ≥ 0.38 for both Jaccard and Bray-Curtis distances) (Figure S1c,d).

Adding additional samples over time yielded similar marginal vertebrate richness as adding additional samples spatially (permutation test *p* = 0.20, Figure 4). Sites that had higher vertebrate richness on day 1 did not have a greater number of OTUs on day 2 (R^2^ = 0.03, p = 0.41) (Figure S2c), nor was there a significant difference in richness between the two days among sites (paired t-test, *p* = 0.55). Running the fans for just 1 day, instead of adding a second day, sampled 19% fewer vertebrate species (128 for the 28 samples for day 1 vs. 157 species for both days) and an asymptotic richness that was 27% lower (150 species for only day 1 samples vs. 206 for both days) (Figure S2d). There was no significant difference in composition by day for presence/absence or relative abundance (PERMANOVA by day, *p* > 0.32; Figure S4).

Exploring the controls on calculated asymptotic richness of vertebrates, decreasing the number of independent PCR replicates combined for a single sample led to lower observed richness and predicted asymptotic richness. At 1 PCR replicate per sample, richness at 56 samples would have averaged 60% less than with all 12 PCR replicates (59 vs. 157 OTUs), and asymptotic richness would have been 50% less (103 vs. 206 OTUs) (Figure 5c). Doubling sequence depth across all samples was not predicted to alter vertebrate richness (157 OTUs). On average, if we had sampled only half as many site locations (i.e., 14 sites) for both days, vertebrate richness would have been 19.1% lower (127 OTUs) and asymptotic richness 21% lower (163 vs. 206 OTUs) (Figure S3).

## Discussion

Metabarcoding of airDNA for terrestrial biodiversity assessment is only beginning to develop, yet the research presented here suggests that it has as much potential as metabarcoding of aquatic eDNA does for aquatic biodiversity. The total richness of arthropods detected with airDNA at Barro Colorado Island (BCI) in this 2-day, 28-site study – 1293 OTUs – is similar to the richness detected with more conventional techniques. For example, when Souto-Vilaros et al.^25^ ran two sets of 10 bucket light traps at BCI in the 2021 dry season and then used metabarcoding on homogenized collections of arthropods, they detected an average of 365 BOLD Barcode Index Numbers (BINs, which are analogous to our 97% OTUs) per trap per day and a total of 2974 BINs across all 20 trap-days. A week of insect collection with a single Malaise trap during the dry season in a seasonally dry forest in Costa Rica could yield 100 - 500 species depending on the location^24^. Compared to other airDNA studies, arthropod richness detected here is higher than in other studies in temperate ecosystems^21,30^.

The number of vertebrates detected with airDNA in the present study is similar to or greater than that often detected using more conventional techniques. Here, one could conservatively estimate that the 157 OTUs that were detected represent approximately 149 species - slightly less than half of the at least 350 vertebrates known on BCI. By comparison, just 19 species of mammals were observed on BCI with 20 camera traps deployed for 1 calendar year^31,32^, which is similar to other camera-trap surveys in neotropical forests^33–35^. Campos-Cerqueira et al.^35^ documented ∼50 bird species on BCI during a four-week deployment of passive acoustic monitoring, but the equivalent of a month’s worth of passive acoustic monitoring recordings in Brazilian forests detected approximately 150 species of birds^36^. Conventional point-count surveys in tropical forests often record over 90 bird species per site^37,38^. Rodgers et al.^15^ sampled the DNA from 1084 carrion flies collected over 29 days in BCI and detected the DNA of 20 mammal, 4 bird, and 1 lizard species. In addition, the vertebrate richness detected at BCI in this study of airDNA exceeds previous richness detected in temperate ecosystems by sampling airDNA^21,22,30^.

The relative abundance of different families of arthropods observed with airDNA differs from distributions obtained with traditional techniques. With airDNA in this study, almost 80% of the COI OTUs detected were Cecidomyiidae (gall midges). In their insect DNA survey on BCI using bucket light traps, Souto-Vilarós et al.^25^ also found that Cecidomyiidae were the dominant family, though they comprised only about 12% of the species detected. Other studies in different regions also report a high abundance of Cecidomyiidae from traditional techniques, but not at the levels detected with airDNA here^21,39,40^. Recent studies that employed DNA barcoding have suggested that Cecidomyiidae is a hyperdiverse insect family with many undescribed taxa^41^, so there is no reason to assume that airDNA is overrepresenting the diversity of Cecidomyiidae on BCI. Cecidomyiidae are small and may be underrepresented with traditional trapping methods, so the proportion of Cecidomyiidae DNA in air might be higher than conventional surveys would estimate. Alternatively, the strong fan system that we used could preferentially capture weak-flying insects such as Cecidomyiidae on the filter, which then overwhelms our ability to detect other arthropod taxa. Given that we detected more vertebrates using a vertebrate-specific 12S rRNA assay than with the general COI primers, it is also likely that more DNA from other arthropod species outside of Cecidomyiidae is present on the filters, but its amplification and sequencing was overwhelmed by the more abundant Cecidomyiidae DNA.

Examining the relative abundance of different bird species, there are some similarities between airDNA and other techniques of assessing bird assemblage composition, but the airDNA assemblage appears to be unique. The bird species with the highest occurrences in the airDNA data (*Coragyps atratus*, *Amazona farinosa*, and *Tinamus major*) are all abundant on the island and are among the most observed during Christmas Bird Counts^42^ and by eBird observers in February and March^43^. Yet, there is little overall correspondence between airDNA and these lists. For example, *Amazona farinosa* is the 2nd most observed species in the Christmas Bird Count, but the most observed species with Christmas Bird Counts, *Psarocolius montezuma*, was not detected with airDNA at all. A large proportion of the birds whose DNA was captured here are not captured with mist nets at all^44^, but the airDNA abundances do not necessarily reflect the relative abundance of bird species caught in nearby mist nets when restricting the analysis to these species.

The composition of mammals in the airDNA assessment was similar to that observed by conventional methods yet was different from the distribution of species known to be present on the island. For example, bats make up approximately half of the mammal species on the island^45^, but only 4 bat species were detected with airDNA. In contrast, there were strong similarities in abundance between airDNA and the abundances of species detected with camera traps. Rowcliffe et al.^31^ lists abundances of 15 species of mammals in the understory as detected with camera traps over a one-year period (2008-9). Nine of the 10 most abundant species were detected with airDNA, and the 3 most observed species with camera traps were among the top 4 detected with airDNA. Meanwhile, *Nasua narica* had the third highest occurrence rate with airDNA, but the seventh most for camera traps.

It appears clear from this work that analysis of airDNA quantifies a high richness of arthropods and vertebrates that can complement more traditional techniques. By sampling 28 field sites across two days, we can also evaluate how sampling strategy affects species detections to help inform future sampling. Lynggaard et al.^22^ found that vertebrate airDNA composition differed across 3 sampling days in a mixed forest in Denmark. We also found a benefit in sampling a second day for each location, as this second day allowed us to detect 50% more unique OTUs for both arthropods and vertebrates. Yet, this marginal gain in richness from a second day of sampling was only 49% of the marginal richness of adding an additional site on the first day, while there was no significant difference in the marginal richness between additional sites or days for vertebrates. As such, these results suggest that sampling designs should maximize the number of independent sites sampled first. As for how best to array airDNA samplers, there was no pattern in composition with distance between any two samples for either arthropods or vertebrates, so presumably airDNA samplers could have been arrayed more closely to one another with no reduction in species detection.

Our experimental design of processing 12 PCR replicates separately for each sample also provides some insights into how to maximize the efficiency of the laboratory work. For example, there appeared to be little benefit to processing more than 6 COI PCR replicates for arthropods to detect maximal OTU richness across sites, but the additional 6 12S rRNA PCR replicates still generated over 20 additional vertebrate OTUs. We observed little benefit of additional sequencing depth for individual PCR replicates, so there would be little reason to recommend greater sequencing depth in the future. In general, increasing both field effort (i.e., the number of samples and the number of days a site was sampled) and laboratory effort (i.e., the number of PCR replicates) increased the estimated asymptotic richness, which follows general patterns of relationships between survey completeness and estimated asymptotic richness when there is species turnover among sites^46–49^. In general, it is unclear what a standard methodology would be most appropriate for airDNA analyses which would allow for robust comparisons of richness across sites and over time. As with most biodiversity assessments, comparing richness using static methodologies can generate biased patterns^46,48,50^.

One thing that is clear is that more work is necessary to improve metabarcoding reference databases for COI, as well as other marker gene regions. Many vertebrate species were only detected when analyzing samples with the 12S primers. It is likely that COI DNA from these species was also present on the filters, but less likely to be detected because of the more abundant arthropod DNA. Unless COI primers can be identified that specifically amplify vertebrates, quantification of vertebrate richness will likely require metabarcoding in other regions besides COI. It is often noted that arthropod COI reference databases need to be improved to reduce uncertainty in taxonomic assignment^51,52^, but so will vertebrate reference databases outside of COI. The COI primers used here amplify a 313-bp region within the Folmer region with the reverse primer at the same location as the reverse Folmer primer, but with degenerate bases to broaden taxon amplification^53^. With Oxford Nanopore sequencing, it is possible that the full Folmer fragment could be amplified, which would allow for more complete matching with the BOLD database. Yet, whether amplifying a longer fragment is possible with airDNA or would lead to biases in DNA detection or taxonomic assignment – and, thus, patterns of diversity – is unknown.

Despite the need to further develop the technique and find standards that will allow for meaningful comparisons across locations and over time, airDNA metabarcoding can soon be operationalized at scale for terrestrial biodiversity assessment. In less than 72 hours, we were able to deploy 28 airDNA samplers over a 1.5 km^2^ area, collect two consecutive 24-hour samples, and then rapidly process the samples in the laboratory and analyze the data to survey faunal biodiversity at levels similar to or better than conventional faunal survey techniques. With this system, the field apparatus is relatively inexpensive and utilizes standard components whose deployment can be scaled with little training. A single commercial lab would be able to process tens to hundreds of thousands of samples per year, providing more than enough capacity for any terrestrial biodiversity surveys that are likely to be generated any time soon.

Raw data and processed results could easily be archived in repositories such as NCBI and the Global Biodiversity Information Facility. Although conventional techniques for biodiversity assessment are largely limited to a single taxonomic group, a single airDNA filter can generate biodiversity data for arthropods and vertebrates. Plants and fungi could easily be added, too^26,54^, creating a single pan-taxa biodiversity survey that could be the key to scaling rapid terrestrial biodiversity assessments, unlocking the broad adoption of conservation monitoring and biodiversity credits.

## Methods

This study was conducted on Barro Colorado Island (BCI), in Lake Gatun in the Republic of Panama. BCI (9.15, -79.85) is a 1542--hectare lowland tropical forest reserve that sits at 85 - 120 m above sea level, is part of the Barro Colorado Nature Monument, and was created in 1913 during the flooding of the Chagres River for the Panama Canal. BCI receives an annual average rainfall of approximately 2600 mm and has an annual average daily maximum air temperature of 28.5 °C. Although historically the land that is now BCI had multiple land uses, the island is now nearly 100% forested. The island has a known mammal community of 108 species^15^, over 150 birds^55^, and at least 100 reptiles and amphibians^56^. However, some of these organisms, such as the jaguar (*Panthera onca*), are infrequent visitors to the island, and others, such as migrating bird species, may be found only at certain times of the year. The arthropod community of BCI is estimated to include over 5000 species, of which more than 2300 have been described taxonomically and recorded as part of long-term monitoring by the Forest Global Earth Observatory (ForestGEO)^25,57,58^.

Twenty-eight airDNA sampling sites were established along 4 trails (4 on Snyder-Molina, 9 on Shannon, 8 on Barbour Lathrop, and 6 on Fairchild) as well as one location near the water in the laboratory area (Figure 6, Table S1). Trail sampling sites were all forested (typically secondary forest <100 years in age), spaced approximately 100 m apart, and located within a few meters from the trail, which typically consisted of no more than evenly spaced cinderblocks sunk to ground level.

**Figure 6.**
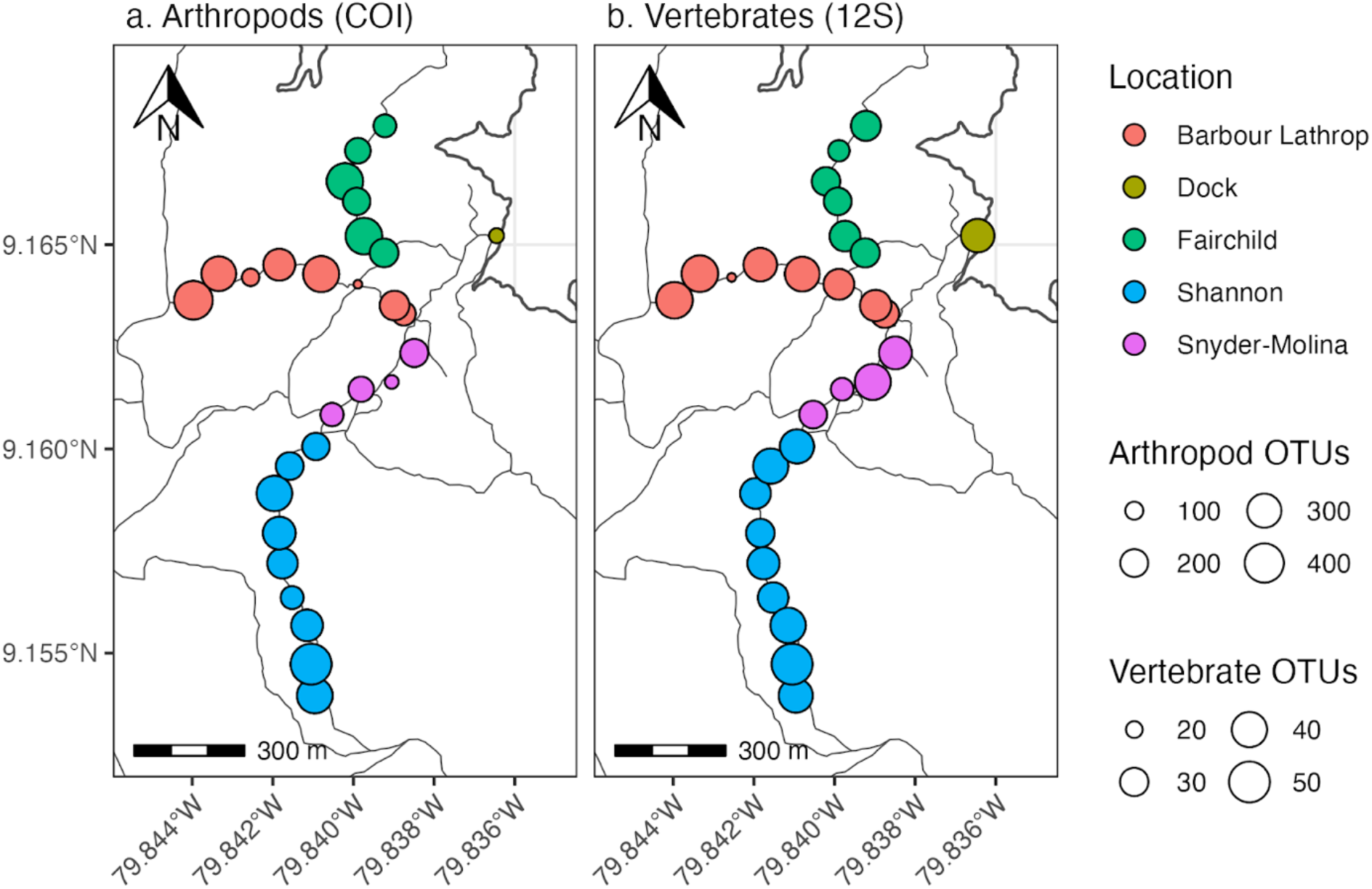
Map of air sampling sites on Barro Colorado Island showing the observed OTU richness of arthropods with the COI primer set (a) and vertebrates from the 12S primer set (b) per sampling location across days 1 and 2. Thin gray lines represent maintained trails.

For air sampling, 0.7A 120-mm fans were connected to custom-designed brackets and then strapped onto trees with 1-inch webbing adjacent to the trail at approximately 1.5 m height (Figure 7). Fans were connected to a 17 amp-hour 12V lead acid battery, a custom air filter was placed in front of the fan, and then a plastic grate attached to secure the filter card in place.

**Figure 7.**
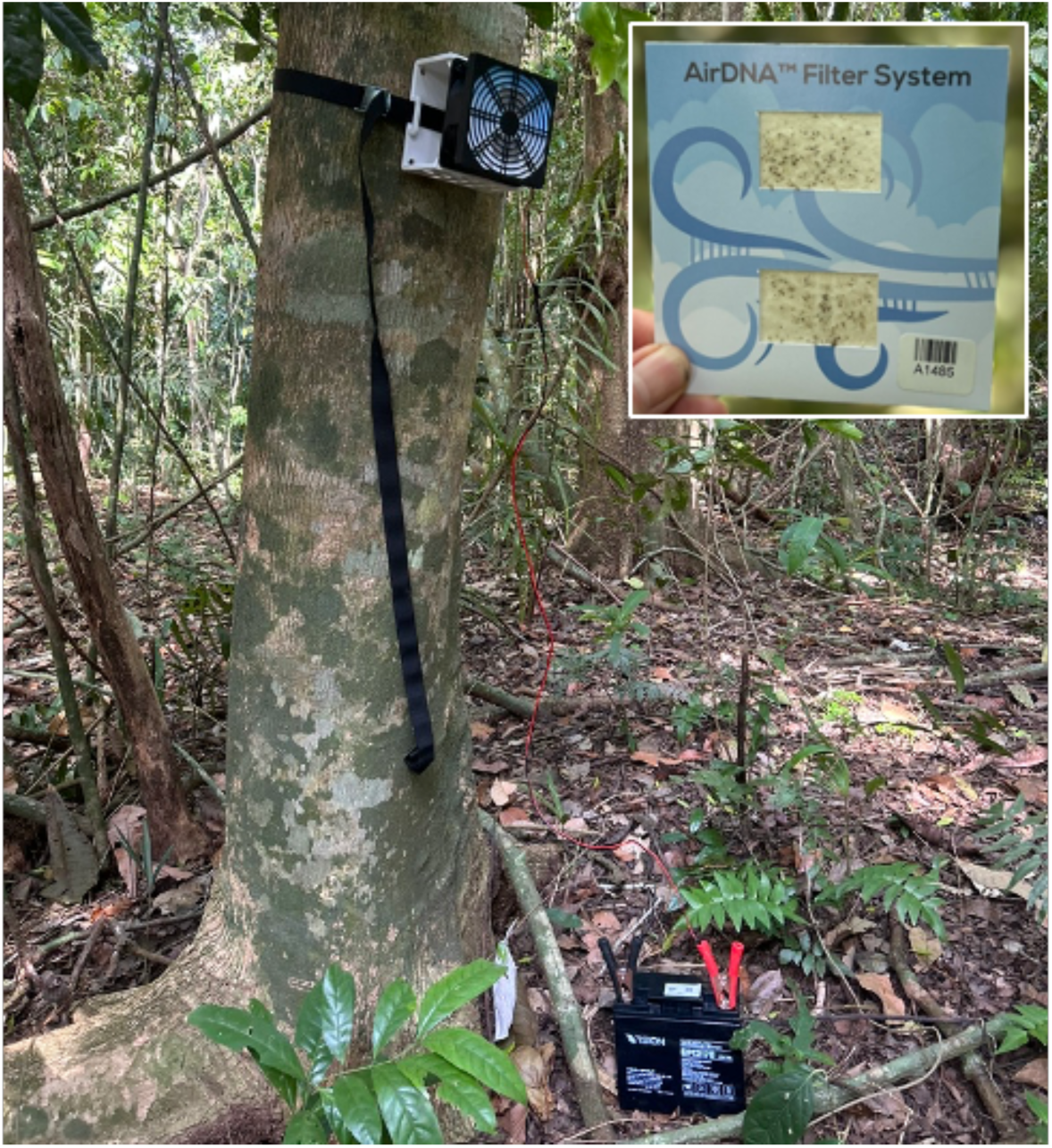
Deployed airDNA sampling system, consisting of a fan attached to a bracket, which is affixed to a tree at eye level and connected to a battery on the ground. Inset: airDNA filter cartridge after running 24 hours.

Filter cards contained 2 10-cm^2^ windows that held polyester filter fabric with a MERV13-rated filter in place. Samples were collected on two consecutive days at each sampling location.

Lab testing showed that the fans pass approximately 216 m^3^ h^-1^ with no filter in place and 36 m^3^ h^-1^ with filters installed. Filtering was initiated between 10:00 and 17:10 on March 11, 2025 (Day 1) and 8:30 and 17:40 on March 12, 2025 (Day 2). On average, air was filtered for 23.5 h (range 21.2 h to 25.1 h). For collection, the filter was removed, added to a polyethylene bag with silica desiccant, and stored at 4 °C later that day. On March 15, all filters were transported at ambient temperature to the Jonah Ventures laboratory in Boulder, CO, USA where they were subsequently frozen before being extracted on March 18.

In the laboratory, DNA was extracted from one of the two windows per filter card, as well as 12 unused filters to serve as blanks. For DNA extraction, we used the Omega Biotek Mag-Bind® Universal Pathogen Core Kit (4x96 Preps) (# M4030-01) according to the manufacturer’s protocol. Extractions were carried out on a Hamilton STARlet running Venus 4, using a method developed by the extraction kit manufacturer and adapted as needed. Approximately 300 µL of lysate was used for genomic DNA extraction. After extraction, DNA was eluted into 100 µL elution buffer and stored at -20 °C.

From each genomic DNA sample, hypervariable regions of the mitochondrial cytochrome c oxidase subunit I (COI) and 12S ribosomal RNA (rRNA) genes were PCR amplified. The COI primers were mlCOIintF (5’-GGWACWGGWTGAACWGTWTAYCCYCC-3’) and jgHCO2198 (5’- TAIACYTCIGGRTGICCRAARAAYCA-3’)^53^. The 12S rRNA primers were 12SVert_F (5’- ACTGGGATTAGATACCCYACTATG-3’) and 12SVert_R (5’-GAGRRTGACGGGCGGTDT-3’) (adapted from Evans et al.^59^ primers 12SF and Ac12S with one and three degenerate nucleotides, respectively). Forward and reverse primers also contained a 5’ adaptor sequence to allow for subsequent indexing and sequencing. For both primer pairs, 12 replicate aliquots of extracted DNA per sample were PCR amplified.

Each 25 µL PCR reaction was mixed according to the Promega PCR Master Mix specifications (Promega # M5133, Madison, WI) which included 12.5 µL Master Mix, 0.5 µL of each 10 µM primer, 2.0 µL of gDNA, and 10.5 µL DNase/RNase-free H2O. DNA was PCR amplified using the following conditions: for COI, initial denaturation at 94 °C for 2 minutes, followed by 45 cycles of 15 seconds at 94 °C, 30 seconds at 50 °C, and 1 minute at 72 °C, and a final elongation at 72 °C for 10 minutes. For 12S rRNA, initial denaturation at 94 °C for 3 minutes, followed by 45 cycles of 30 seconds at 94 °C, 30 seconds at 52 °C, and 1 minute at 72 °C, and a final elongation at 72 °C for 10 minutes. Amplicons were then cleaned by incubating the reactions with Exo1/SAP (New England Biolabs # M0293L and # M0371L) for 30 minutes at 37 °C followed by inactivation at 95 °C for 5 minutes and storage at -20 °C.

A second round of PCR was performed to complete the sequencing library constructs and integrate dual 10-bp multiplexing index sequences. The indexing PCR included Promega Master mix, 0.5 µM of each primer and 2 µL of template DNA (cleaned amplicon from the first PCR reaction) and consisted of an initial denaturation of 95 °C for 3 minutes followed by 8 cycles of 95 °C for 30 sec, 55 °C for 30 seconds, and 72 °C for 30 seconds.

Final indexed amplicon pools from each sample were cleaned and normalized using a magnetic bead-based protocol based on Buchner^60^ with minor alterations. A 15-µL aliquot of PCR amplicon was purified and normalized using Cytiva SpeedBead magnetic carboxylate modified particles (# 45152105050250). Samples were then combined into a single multiplexed library by adding 5 µL of each normalized sample to the pool.

Library pools were prepared for sequencing on an Oxford Nanopore Technologies MinION flow cell (Oxford, England) using the Ligation Sequencing Kit V14 (# SQK-LSK114) before being run on a single flowcell on a GridION sequencer. For the COI library, the sequencing run was stopped after approximately 48 hours, at which point 30.6M reads had accumulated and pore availability was low. For the 12S rRNA library, sequencing ended after 72 hours and 21.6M reads had accumulated.

Raw Nanopore sequencing output was converted from pod5 to fastq format using Dorado v0.7.1 (Oxford Nanopore Technologies 2024) and the super-high accuracy basecalling model (dna_r10.4.1_e8.2_400bps_sup@v5.0.0). Cutadapt v3.4^61^ was then used to trim outer adapters and reorient reads in a consistent 5’ to 3’ direction. The resulting reads were sorted into individual samples by demultiplexing with pheniqs v2.1.0^62^, allowing no more than one mismatch in each of the paired 10 bp indices. Cutadapt was then used again to extract the target amplicon by removing the gene primers, discarding any reads where one or both primers were not found or where the resulting amplicon sequences were < 298 bp or > 328 bp for COI and < 300 or > 450 for 12S rRNA. Exact sequence variants (ESVs) were then identified for each sample using the unoise3 denoising algorithm^63^ as implemented in vsearch^64^. Only reads that were observed 4 or more times across all samples and had a maximum expected error rate < 1 bp^65^ were considered as candidate ESVs and putative chimeras were filtered using the uchime3 algorithm^63^. Final read counts were determined for each sample by mapping unfiltered raw reads to the identified ESVs using the vsearch function usearch_global with a minimum identity of 95%. For each final ESV, a consensus taxonomy was assigned using a custom best-hits algorithm and a reference database consisting of publicly available sequences on NCBI GenBank^66^, Barcode of Life Data System (BOLD)^67^, and Jonah Ventures voucher sequence records. Reference database searching used vsearch to conduct exhaustive semi-global pairwise alignments, and final match quality was quantified using a query-centric approach, where the percent match ignores terminal gaps in the target sequence, but not the query sequence. A consensus taxonomic assignment was then generated using either all 100% matching reference sequences or all reference sequences within 1% of the top match, accepting the reference taxonomy for any taxonomic level with >90% agreement across the top hits. We retained only ESVs that matched at 85% or higher to arthropods or vertebrates in the NCBI or BOLD reference databases. All human and domestic animal sequences (e.g. dog, chicken, cow) were removed. A total of 4.9M reads were analyzed after filtering for COI and 1.2M reads for 12S.

ESVs were then curated by occurrence and abundance using LULU^68^, with a minimum parent:child abundance ratio of 10:1 to remove spurious sequences. We then clustered LULU-curated ESVs with 97% identity thresholds to generate OTUs that would more closely align with species. Although there are examples of species that have an interspecific distance less than 3% divergence^41^, we conservatively used a final 97% clustering distance for both COI and 12S^25,69^. We retained only ESVs that matched at 85% or higher to arthropods or vertebrates in the NCBI or BOLD reference databases.

All statistical analyses were performed in R v.4.3.0^70^. To estimate the total observed and predicted species pools across PCR replicates and samples, observed OTU accumulation was calculated from incidence data and then extrapolated to asymptotic richness using ‘estimateD’ (yielding Chao2 estimates) and ‘iNEXT’ in iNEXT v.2.0.20^71^. To explore compositional differences across geographic distance and days, we used non-metric multidimensional scaling (NMDS) ordinations of pairwise presence/absence Jaccard similarities and relative abundance-based Bray-Curtis similarity using ‘metaMDS’ in vegan^72^. Visualizations were rendered with ggplot2^73^, ggspatial^74^, and treemapify^75^. For COI, only arthropod sequences were analyzed statistically, and the vertebrate sequences were summarized qualitatively.

## Acknowledgments

The authors acknowledge the assistance of the Josh Tewksbury and BCI staff in helping facilitate this research. Lovisa Duck and Adalberto Gomez provided invaluable assistance in the field.

## Author contributions

JC and NF conceived of the research and carried out the field component of the research. JD was responsible for the laboratory analyses, DL for the bioinformatic processing, and NS and JC for the data analyses. All authors contributed to the writing.

## Additional Information

Jonah Ventures is a private commercial entity.

## Data availability

The raw sequencing data generated in this study have been deposited in the NCBI Sequence Read Archive (SRA) under BioProject accession number **PRJNA1282780**. The data can be accessed at https://www.ncbi.nlm.nih.gov/bioproject/PRJNA1282780.

## Supplementary Materials

Text S1. Description of vertebrate COI ESVs.

Only 14 COI ESVs were assigned to birds and mammals. We detected the DNA of 9 bird species: snail kite (*Rostrhamus sociabilis*, 100%), black vulture (*Coragyps atratus*, 100% match), grey-chested dove (*Leptotila cassinii*, 100%), greater ani (*Crotophaga major*, 100%), bay-breasted warbler (*Setophaga castanea*, 100%), black-crowned antshrike (Thamnophilus atrinucha, 100%), red-lored amazon (Amazona autumnalis, 100%), blue-headed parrot (Pionus menstruus, 100%), and great tinamou (Tinamus major, 99%). Five mammal species were also detected: collared peccary (Dicotyles tajacu, 98%), white-nosed coati (Nasua narica, 91% match), mantled howler monkey (Alouatta palliata, 100%), three-toed sloth (Bradypus variegatus, 100%), and lowland paca (Cuniculus paca, 96%). 6 of the 9 bird species found with COI were also found unambiguously with 12S rRNA. *Leptotila cassinii, Amazona autumnalis,* and *Setophaga castanea*, which were identified with the COI assay, were not identified unambiguously with 12S, but each of these taxa was a possible hit for OTUs with ambiguous species-level taxonomy.

**Table S1.** Site locations and 97% OTU richness per sample (site-by-day) and amplicon (arthropod = COI, vertebrate = 12S rRNA).

| Site | Location | Latitude | Longitude | Sample (Day 1) | Sample (Day 2) | Arthropod OTUs (Day 1) | Arthropod OTUs (Day 2) | Vertebrate OTUs (Day 1) | Vertebrate OTUs (Day 2) |
| --- | --- | --- | --- | --- | --- | --- | --- | --- | --- |
| BL0 | Barbour Lathrop | 9.16331 | -79.83875 | A1476 | A1504 | 97 | 64 | 16 | 22 |
| BL1 | Barbour Lathrop | 9.16351 | -79.83898 | A1477 | A1508 | 104 | 172 | 22 | 21 |
| BL2 | Barbour Lathrop | 9.16403 | -79.83989 | A1478 | A1511 | 48 | 44 | 21 | 19 |
| BL3 | Barbour Lathrop | 9.16428 | -79.8408 | A1479 | A1510 | 234 | 163 | 26 | 22 |
| BL4 | Barbour Lathrop | 9.16451 | -79.84184 | A1480 | A1509 | 170 | 180 | 25 | 24 |
| BL5 | Barbour Lathrop | 9.1642 | -79.84255 | A1481 | A1507 | 16 | 89 | 2 | 17 |
| BL6 | Barbour Lathrop | 9.16429 | -79.84334 | A1482 | A1506 | 257 | 159 | 22 | 28 |
| BL7 | Barbour Lathrop | 9.16364 | -79.84397 | A1483 | A1505 | 316 | 187 | 34 | 19 |
| DF1 | Fairchild | 9.1648 | -79.83924 | A1502 | A1529 | 115 | 129 | 17 | 21 |
| DF2 | Fairchild | 9.16521 | -79.83974 | A1501 | A1528 | 267 | 239 | 25 | 18 |
| DF3 | Fairchild | 9.16606 | -79.83992 | A1500 | A1527 | 155 | 76 | 17 | 21 |
| DF4 | Fairchild | 9.16655 | -79.84021 | A1498 | A1526 | 224 | 234 | 16 | 22 |
| DF5 | Fairchild | 9.1673 | -79.83989 | A1497 | A1525 | 106 | 121 | 12 | 15 |
| DF6 | Fairchild | 9.16791 | -79.83922 | A1496 | A1524 | 122 | 45 | 23 | 18 |
| Dock | Dock | 9.16522 | -79.83645 | A1475 | A1503 | 48 | 49 | 20 | 26 |
| RCS1 | Shannon | 9.16006 | -79.84093 | A1491 | A1519 | 129 | 78 | 31 | 18 |
| RCS2 | Shannon | 9.15958 | -79.84158 | A1499 | A1518 | 124 | 108 | 30 | 20 |
| RCS3 | Shannon | 9.15891 | -79.84196 | A1490 | A1517 | 206 | 212 | 23 | 21 |
| RCS4 | Shannon | 9.15794 | -79.84184 | A1489 | A1516 | 171 | 171 | 23 | 15 |
| RCS5 | Shannon | 9.1572 | -79.84176 | A1488 | A1515 | 162 | 153 | 17 | 27 |
| RCS6 | Shannon | 9.15636 | -79.84152 | A1487 | A1514 | 74 | 95 | 24 | 20 |
| RCS7 | Shannon | 9.15568 | -79.84115 | A1486 | A1549 | 166 | 165 | 27 | 28 |
| RCS8 | Shannon | 9.15473 | -79.84105 | A1485 | A1513 | 297 | 261 | 49 | 16 |
| RCS9 | Shannon | 9.15396 | -79.84096 | A1484 | A1512 | 186 | 223 | 24 | 26 |
| SM1 | Snyder-Molina | 9.16235 | -79.83849 | A1495 | A1523 | 126 | 99 | 22 | 27 |
| SM2 | Snyder-Molina | 9.16164 | -79.83905 | A1494 | A1522 | 39 | 49 | 19 | 33 |
| SM3 | Snyder-Molina | 9.16146 | -79.83981 | A1493 | A1521 | 126 | 83 | 11 | 17 |
| SM4 | Snyder-Molina | 9.16084 | -79.84053 | A1492 | A1520 | 39 | 120 | 23 | 13 |

Tables S2-S5 available in file “BCI Jonah AirDNA March.TablesS2-S5.xlsx”

Table S2. Consensus taxonomy and read counts for each COI OTU and sample PCR replicate.

Table S3. Reference database hits for each COI OTU.

Table S4. Consensus taxonomy and read counts for each 12S OTU and sample PCR replicate.

Table S5. Reference database hits for each 12S rRNA OTU.

**Table S6.** Analysis costs of different sampling and laboratory efforts, using Jonah Ventures 2025 pricing. All costs are in USD.

|  | Number of samples | Filter | Base | Number of assay 1 PCR reps | Cost per assay 1 PCR rep | Cost for 2nd assay (1 rep incl.) | Number of assay 2 PCR reps | Cost per assay 2 PCR rep | Total |
| --- | --- | --- | --- | --- | --- | --- | --- | --- | --- |
| 56 samples, 12 independent PCR reps, 2 assays | 56 | \$5 | \$90 | 11 | \$65 | \$65 | 11 | \$65 | \$89,040 |
| 56 samples, 12 pooled PCR reps, 2 assays | 56 | \$5 | \$90 | 11 | \$5 | \$65 | 11 | \$5 | \$15,120 |
| 14 samples, 6 pooled PCR reps for arthropods | 14 | \$5 | \$90 | 5 | \$5 | | | | \$1,680 |
| 14 samples, 12 pooled PCR reps for vertebrates | 14 | \$5 | \$90 | 11 | \$5 | | | | \$2,100 |
| 14 samples, 6 pooled PCR reps for arthropods, 12 pooled PCR reps for vertebrates, 2 assays | 14 | \$5 | \$90 | 5 | \$5 | \$65 | 11 | \$5 | \$3,360 |

**Figure S1.**
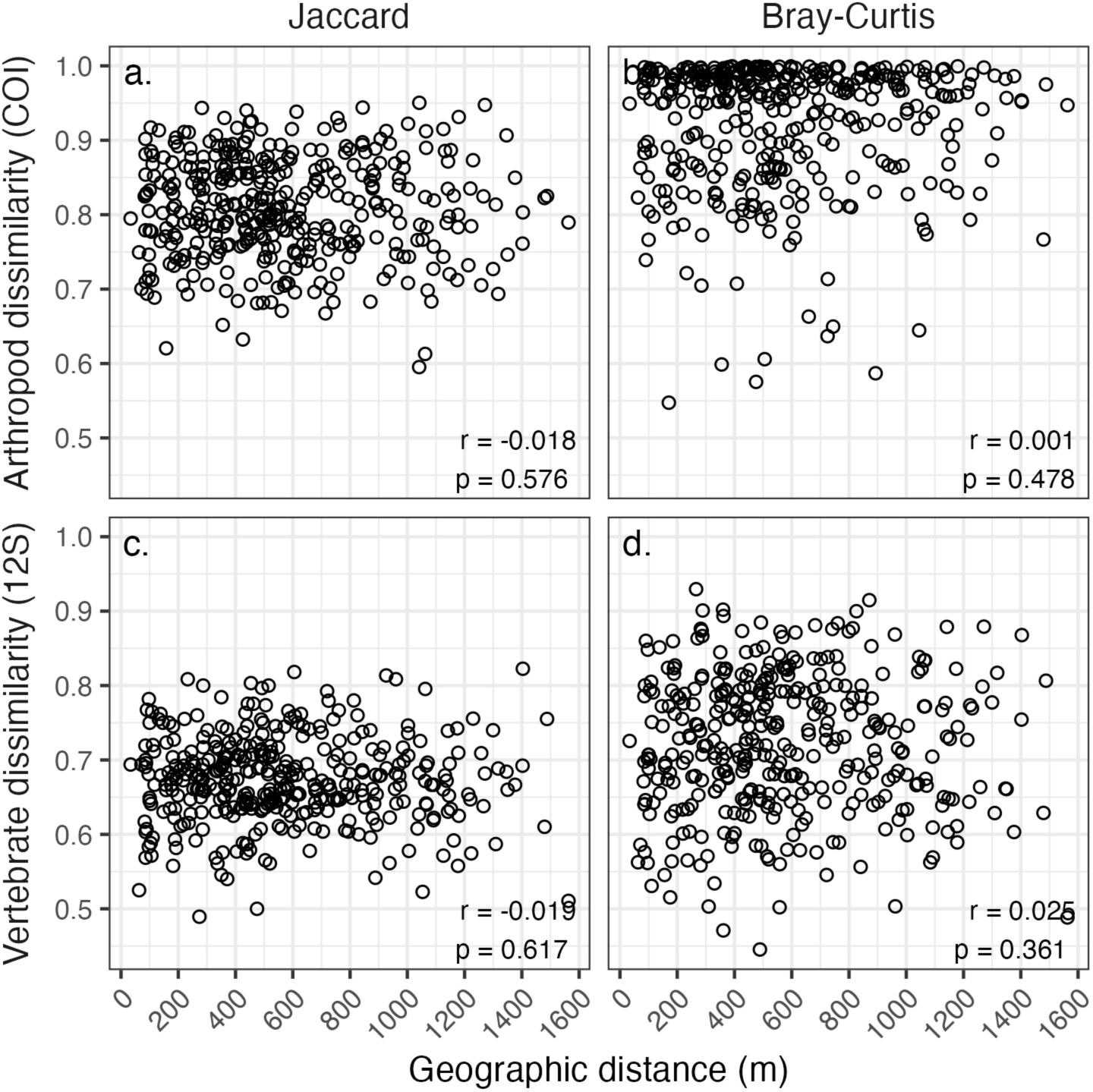
Relationships between pairwise geographic distance of sites and pairwise assemblage dissimilarity of sites (day 1 and 2 sample reads summed for each site) for (a,c) presence/absence (Jaccard) and (b,d) relative abundance (Bray-Curtis) of (a,b) COI arthropods and (c,d) 12S vertebrates. Reported coefficients and p-values are from Mantel tests using Spearman rank correlations and permutation-based significance testing.

**Figure S2.**
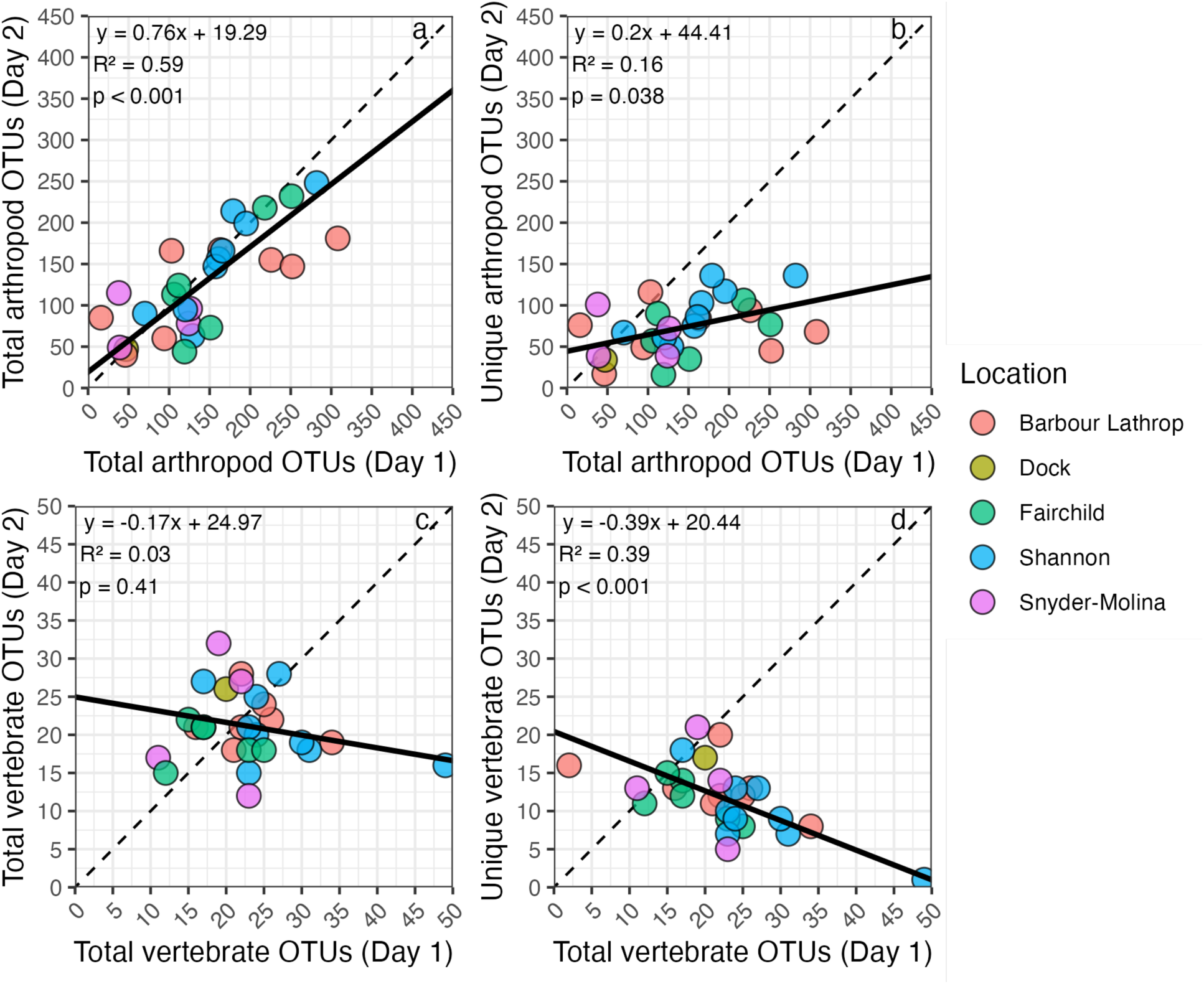
Comparison of day 1 and day 2 (a,c) total OTU richness by day per site and (b,d) total OTU richness for day 1 and marginal (unique) OTU richness for day 2 per site for (a,b) COI arthropods and (c,d) 12S vertebrates. “Unique” refers to OTUs that were not present at a given site for day 1, but might have been present in a different site on day 1 or 2. Solid lines, equations, R² values, and p-values represent reduced major axis (RMA) regression results.

**Figure S3.**
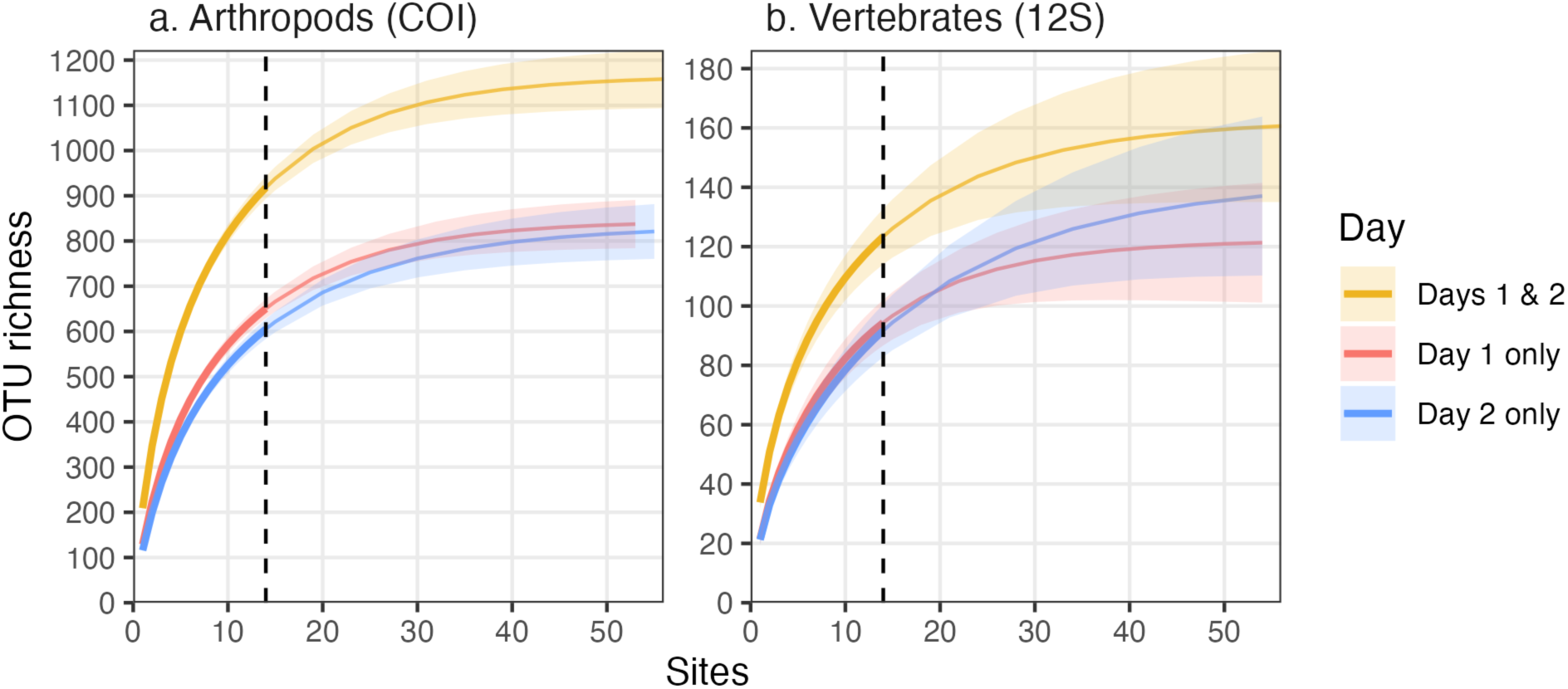
Accumulation of OTUs by adding up to 14 sampling sites (randomly drawn 10 times) for (a) arthropods and (b) vertebrates across samples after summing PCR replicate reads per sample. For Days 1 & 2, OTU reads were summed by site across days. Vertical dashed lines represent the number of sites (n = 14), and dashed segments of curves represent extrapolated accumulation to 4x the number of sites.

**Figure S4.**
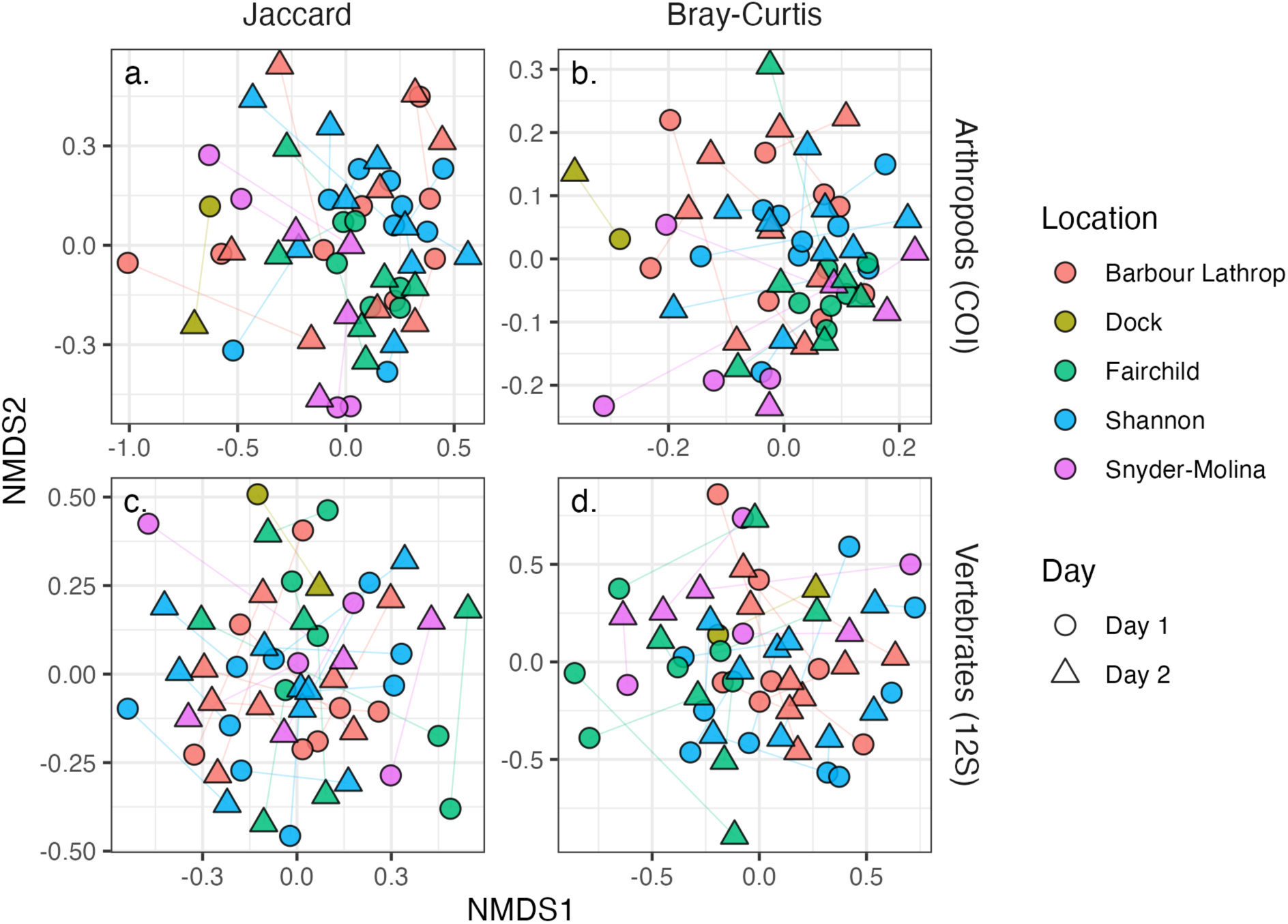
Non-metric multidimensional scaling ordinations of (a,c) Jaccard presence/absence and (b,d) Bray-Curtis relative abundance for (a,b) arthropods and (c,d) vertebrates. Lines connect day 1 and day 2 assemblages for the same site.

